# Complete and validated genomes from a metagenome

**DOI:** 10.1101/2020.04.08.032540

**Authors:** Daniel J Giguere, Alexander T Bahcheli, Benjamin R Joris, Julie M Paulssen, Lisa M Gieg, Martin W Flatley, Gregory B Gloor

## Abstract

The assembly and binning of metagenomically-assembled genomes (MAGs) using Illumina sequencing has improved the genomic characterization of unculturable communities. However, short-read-only metagenomic assemblies rarely result in completed genomes because of the difficulty assembling repetitive regions. Here, we present a strategy to complete and validate multiple MAGs from a bacterial community using a combination of short and ultra long reads (N50 > 25 kb). Our strategy is to perform an initial long read-only metagenomic assembly using metaFlye, followed by multiple rounds of polishing using both long and short reads. To validate the genomes, we verified that longs reads spanned the regions that were not supported by uniquely mapped paired-end Illumina sequences. We obtained multiple complete genomes from a naphthenic acid-degrading community, including one from the recently proposed Candidate Phyla Radiation. The majority of the population is represented by the assembled genomes; recruiting 63.77 % of Nanopore reads, and 64.38 % of Illumina reads. The pipeline we developed will enable researchers to validate genomes from metagenomic assemblies, increasing the quality of metagenomically assembled genomes through additional scrutiny.

## 0.2 Introduction

Metagenomic sequencing and assembly has vastly improved our ability to discover novel genes and pathways from unculturable organisms. Recently, metagenomic sequencing was a key methodology in distinguishing a new branch from the tree of life (Hug et al. 2016). Current methods for generating metagenomically assembled genomes (MAGs) typically combine metagenomic assembly tools, such as Megahit and metaSpades, (Li et al. 2015; Nurk et al. 2017) with binning algorithms such as CONCOCT, BinSanity, and dasBin (Alneberg et al. 2014; Graham, Heidelberg, and Tully 2017; Sieber et al. 2018) to generate a collection of contiguous DNA sequences with similar properties, where each bin may represent a single MAG. Generating and refining bins can be accomplished with tools such as Anvi’o (Eren et al. 2015) and can produce many MAGs from a single sample. The quality of each MAG is typically assessed by several criteria (minimum information proposed here (Bowers et al. 2017)) such as the number of contigs, the percentage of expected single copy core genes present (Campbell et al. 2013), the redundancy of these genes, the number of 23S, 16S, and 5S rRNA genes, and tRNA genes. Generating complete genomes from metagenomes is often difficult or impossible with Illumina-only sequencing because of gaps, local assembly errors, and contamination by fragments from other genomes. Of the thousands of MAGs that have been deposited to public databases, only approximately 60-70 have been completed (Chen et al. 2020), many of these belonging to Candidate Phyla Radiation due to their smaller genome size.

A typical method for assessing the quality of assembled isolate genomes is mapping paired-end Illumina reads against the assembly and evaluating the number of mis-assemblies using a tool such as REAPR (Hunt et al. 2013). Assuming full coverage of the genome with Illumina reads, REAPR is effective at identifying mis-assemblies and validating assemblies for isolates. However, the implicit assumptions of REAPR is that each read pair maps uniquely to the genome, and the Illumina reads are equally distributed. These assumptions may not hold true in metagenomic datasets due to conserved or duplicated genes in the population (16S rRNA genes, transposases, etc.). Although other tools such as Quast (Gurevich et al. 2013) are available, the statistics they provide either require a reference genome, or are only informative for incomplete genome assemblies (such as N50, % identity to reference, etc.). These tools may be useful for Illumina-only assemblies, as well as assemblies with a known reference, however, to the best of our knowledge, a method or pipeline to evaluate genome assemblies without a reference using both short and long reads does not yet exist.

A major issue with completing MAGs with short reads only is that repetitive regions (such duplicated genes, inverted repeats, etc.) are difficult to assemble through. Recently developed technologies such the Oxford Nanopore minION platform can produce reads that span the entire length of repetitive regions, and provide genomic context by anchoring to adjacent regions. While the consensus sequence for these regions will not have a quality score (Q-score) as high as the Illumina supported regions, the Q-scores and coverage may still be high enough to support contiguity with a reasonably acceptable Q-score. Recent benchmarking studies have shown that average Q-scores of 30 are achievable with Oxford Nanopore (Ryan R Wick, Judd, and Holt 2019), and this will likely improve with new algorithmic and technological advances in the future.

There is currently no consensus in the literature on whether the best approach for hybrid genome assembly is by performing short read assembly following by scaffolding with long reads, or by correcting long read assembly with the more accurate short reads. Long reads generated by the Oxford Nanopore Technologies minION platform have been incorporated in recently published hybrid assemblers (Ryan R Wick et al. 2017; Bertrand et al. 2019) to scaffold short read assemblies. These tools can be successful at generating circularized genomes from isolate data, however, Unicycler is explicitly not designed for metagenomic data, and while OPERA-MS improves the contiguity of MAGs, our experience is that it will typically will not generate fully circularized MAGs directly from a high-coverage metagenomic datasets. A recently published alternative is to use a long-read-only metagenomic assembler, metaFlye (Kolmogorov et al. 2019). A potential issue with long-read only assemblies is the high per-base error rate due to the lower quality reads produced by Oxford Nanopore platforms (typical runs can now achieve Q > 10 in high accuracy basecalling mode). To compensate, various polishing algorithms have been proposed to correct remaining single nucleotide polymorphisms (SNPs) or indels, such as Medaka (Oxford Nanopore), Racon (Vaser et al. 2017), and Pilon (Walker et al. 2014). metaFlye has been shown to improve contiguity compared to hybrid assemblers (Kolmogorov et al. 2019). However, many studies using the metaFlye tool have been limited to datasets with relatively “short” long-reads. The metaFlye paper demonstrated improved contiguity using samples from the rumen microbiome (Stewart et al. 2019) and was able to achieve circularized genomes with the Ultra deep sequencing (200X +) of bacterial genomes from the Zymobiomics Mock Community (Nicholls et al. 2019), which have approximate read N50’s of 9kb and 5kb, respectively.

Recent studies have been successful at improving contiguity using long reads (Caceres et al. 2019), and some groups have been reported completing more than a single genome (Stewart et al. 2019; Moss, Maghini, and Bhatt 2020) using Nanopore sequencing. However, there is still a very small percentage of MAGs that have been reported in the literature as complete (~60-70 from 7000 submitted MAGs) (Chen et al. 2020). Chen et al., 2020 also proposed a general workflow for validating circularized genomes. While circularized genomes can now be generated, there are currently no pipelines available to validate these assemblies using long and short reads.

Here, we generate and validate bacterial circularized MAGs from an enrichment culture using metaFlye with ultra-long reads (read N50 > 25 kb) as input, followed by polishing using both long and short reads. To validate each genome, we found that long-reads needed to be strictly filtered by read length, query coverage, and alignment score to avoid mis-mappings. Filtering the long reads allowed us to visualize variants in several genomes where an insertion or deletion was present in a subset of the population. Ten of the 13 initially circulared genomes passed our validation, with coverage as low as 17X. metaFlye is highly succesful in generating many high quality circularized MAGs directly from metagenomic data, however, we demonstrate that additional validation is required to ensure genomic variants are not missed, and that genomes with very low coverage require visual inspection before concluding they are complete.

## 0.3 Methods

### 0.3.1 Sequencing

An algal-bacterial culture initially enriched in 2012 from a a northern Alberta oil sands tailings pond was used for this study. The culture had been routinely propagated since that time. For this study, a 5 mL aliquot was grown in Bold’s Basal media, pelleted, and stored at −80° C until DNA was extracted (Paulssen and Gieg 2019).

DNA extraction was performed to maximize read length (performed with wide-bore pipette tips, slow pipetting, only slow end-over-end inversions). Pelleted cells were resuspended in buffer (50 mM Tris-HCl pH 8.0, 10 mM sodium EDTA, pH 8.0, RNAse A, hemicellulase, lysozyme, zymolyase) and incubated at 37° C for 1 hour, mixing by inversion every 10 minutes. Cetrimonium bromide was added to 2% and NaCl to 1.5 M. The sample was incubated at 50° C for 1 hour, mixing by inversion every 15 minutes. The sample was centrifuged at 6000 x G for 3 minutes. The supernatant was collected and transferred to a new tube. One volume of 25:24:1 phenol:chloroform:isoamyl alcohol pH 8.0 was added, and mixed by inversion. Phases were seperated by centrifugation at 8000 X g for 3 minutes. The aqueous phase was transferred to a new tube, where 1 volume of chloroform was added and mixed by inversion. The phases were separated by centrifugation at 6000 X g for 3 minutes, and the aqueous phase transferred to a new tube. One quarter volume of Tris-EDTA pH 8.0 was added to the chloroform, mixed, and centrifuged as previously. The aqueous phase was removed and combined, and the chloroform extraction was repeated once. After collecting the aqueous phase, sodium acetate (pH 5.2) was added to 0.3 M, and 2 volumes of cold 70% ethanol was added, mixed by inversion. The mixture was centrifuged at 16 000 X G for 2 minutes, and washed once with cold 70% ethanol. The pellet was resuspended in Tris-EDTA pH 8.0, and stored at 4° C until further use. Short DNA fragments were then selectively removed using the Short Read Eliminator (SRE) kit from Circulomics (Baltimore), and the sample was stored at 4° C until sequencing. DNA from the same extraction was used for sequencing on both the Oxford Nanopore minION and Illumina NextSeq 550 platforms.

An Oxford Nanopore minION flow cell R9.4.1 was used with the SQK-LSK109 Kit according to the manufacterer’s protocol version GDE_9063_v109_revK_14Aug2019, with one alteration: for DNA repair and end-prep, the reaction mixture was incubated for 15 minutes at 20° C and 15 minutes at 65° C. Basecalling was performed after the run with Guppy (Version 3.3.0). NanoPlot (De Coster et al. 2018) was used to generate Q-score versus length plots and summary statistics. The read N50 of the unfiltered reads was approximately 24 kb. Nanopore reads were not filtered prior to assembly (as expected by the assembler).

For Illumina sequencing, the Nextera XT kit was used to prepare 2×75 paired-end mid-output NextSeq 550 run at the London Regional Genomics Center (lrgc.ca). Reads were trimmed using Trimmomatic v0.36 (Bolger, Lohse, and Usadel 2014) in paired end mode with the following settings: AVGQUAL:30 CROP:75 SLIDINGWINDOW:4:25 MINLEN:50 TRAILING:15. SLIDINGWINDOW AND TRAILING were added to remove poor quality base calls. Only paired end reads were retained.

### 0.3.2 Assembly

The raw long reads were used for long-read metagenomic assembly using Flye v2.6 (Kolmogorov et al. 2019) with the parameters –meta -g 5m. Circularized contigs larger than 300 kb were extracted as genomes. 300 kb was chosen as a cutoff to remove a circularized mitochondrial genome of an algae that was known to be present in the sample.

### 0.3.3 Extracting correctly mapped Nanopore reads

When mapping Nanopore reads back to the assembly with minimap2 with the parameters –no-secondary (Li 2018), we observed that mis-mapping of nanopore reads occured at a high frequency in the raw output due to the partial mappings, supplementary mappings, and multi-mappings. It was therefore necessary to filter the alignments to ensure only the retention of correctly mapped alignments. We found that filtering long reads with the following criteria (similar to (Caceres et al. 2019)) achieved minimal loss of true mapping, while removing almost all of the mismapped reads.

**Criteria to retain a Nanopore alignment**

- the mapped read is a primary alignment
- the mapped read is > 1000 in length
- the mapped read must have > 90% query coverage
- the ratio of read length to alignment score is < 0.9

Long read length and high query coverage was needed to ensure correct genome location. Furthermore, we found the ratio of read length to alignment score helped remove chimeric and other incorrectly mapped reads (Figure 2). Removing reads with a length divided by score below 0.9 (i.e. highest possible alignment score for a given read length) ensures only the highest quality reads are retained for polishing.

### 0.3.4 Polishing

Initial polishing was done using Rebaler v0.2.0 (R R Wick 2017), which performs multiple rounds of Racon (Vaser et al. 2017) long read polishing. We believe the additional “pruning” step in Rebaler is appropriate for this dataset since it is known that Flye adds a few bases at the point of circularization (Ryan R Wick and Holt 2019). A subsequent round of Pilon v1.23 with the parameters –fix snps,indels –threads 20 –verbose –mindepth 30 –nanopore) (Walker et al. 2014) polishing was performed using Illumina and Nanopore reads. The output from Pilon was retained for validation.

A final round of SNP polishing was manually performed using bcftools v1.10.2 because it was observed that many small indels were not adequately corrected. Using bcftools filter, we retained variants where 50% of the reads support the reference with the parameters -i $MAX(IMF)>0.5. There existed a subset of Illumina reads that “appeared” to not support the insertion because the supporting reads did not span a homopolymer sequence, artificially decreasing the reported IMF value. A cutoff of 50% allowed for most of these regions to be called accurately while minimizing false positives.

### 0.3.5 Validation

Paired Illumina reads were mapped back to the polished MAGs using bowtie2 with the parameters –no-unal –no-discordant –end-to-end –very-sensitive (Langmead and Salzberg 2012), while filtered Nanopore reads were mapped with minimap2 with the parameters -aLQx map-ont –sam-hit-only –secondary=no (Li 2018). Coverage plots were manually generated using the R package circlize (Gu et al. 2014).

## 0.4 Results

### Summary of genomes

The general workflow is shown in Figure 1. Initial assembly with metaFlye generated 13 circularized MAGs. Using a DNA extraction method optimized for high molecular weight DNA is critical; the same community was sequenced and assembled using a bead beating extraction method, achieving a read N50 of about 4.5 Kb, which resulted in only 3 circularized MAGs. After long-read metagenomic assembly with Flye, “pseudo-isolate” subsets of reads for each circularized contig were generated by retaining only the reads that passed the four criteria described in the methods. We found that a distinct cluster of reads with a read length to alignment score ratio from 1-2, contained many chimeric reads (Figure 2).

**Figure 1:**
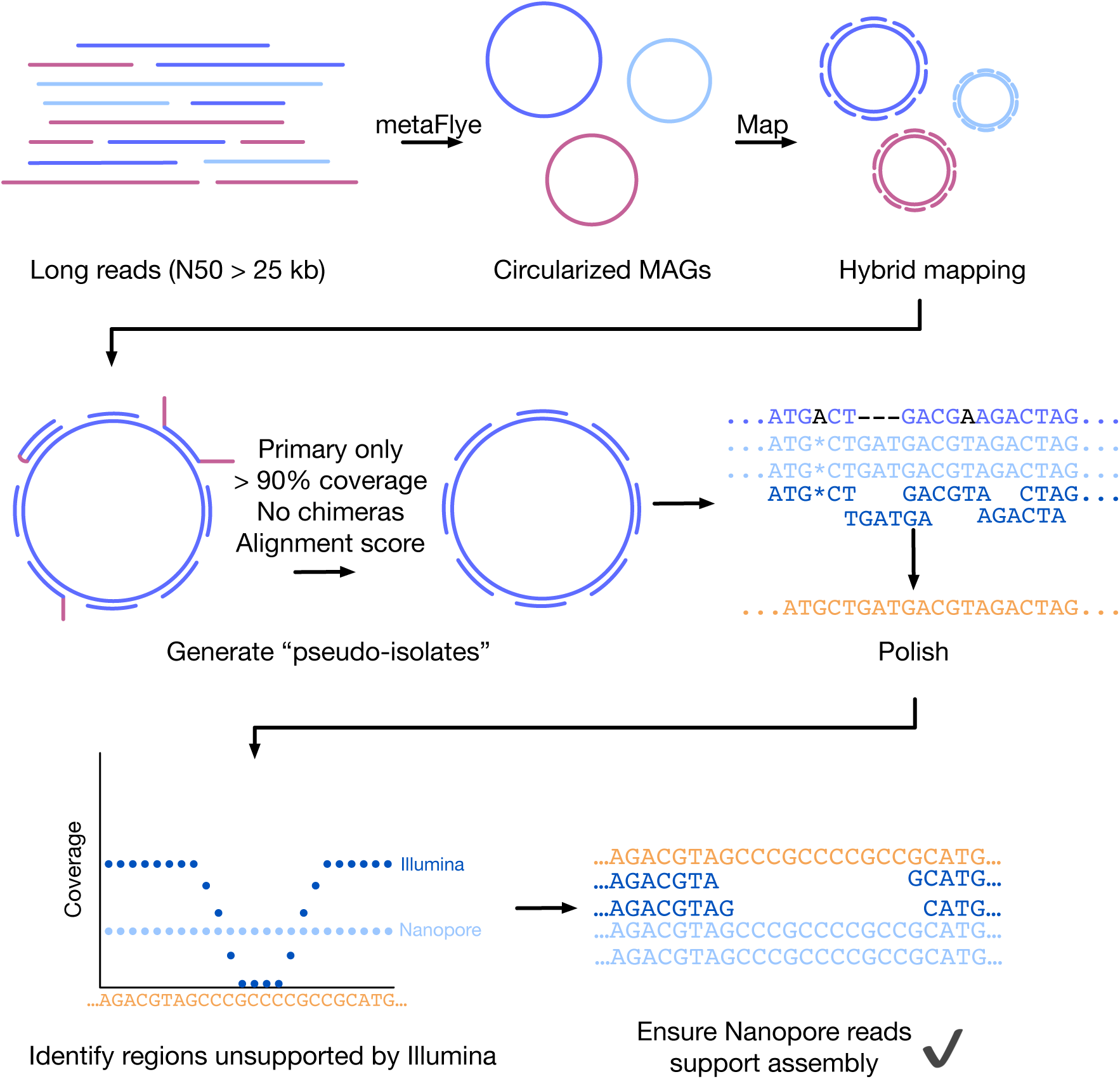
Assembly and validation workflow. Long reads were assembled with metaFlye to generate circularized MAGs, which are then polished with long and short reads (using Rebaler and Pilon, respectively). Pseudo-isolate subsets of reads are generated, that is, subsets of long reads that map to only a single genome. After mapping reads back to each assembly, visualization is performed to find regions that have low coverage support. If the region has sufficient coverage to generate a consensus with long reads only, the long read consensus is used to support the assembly.

**Figure 2:**
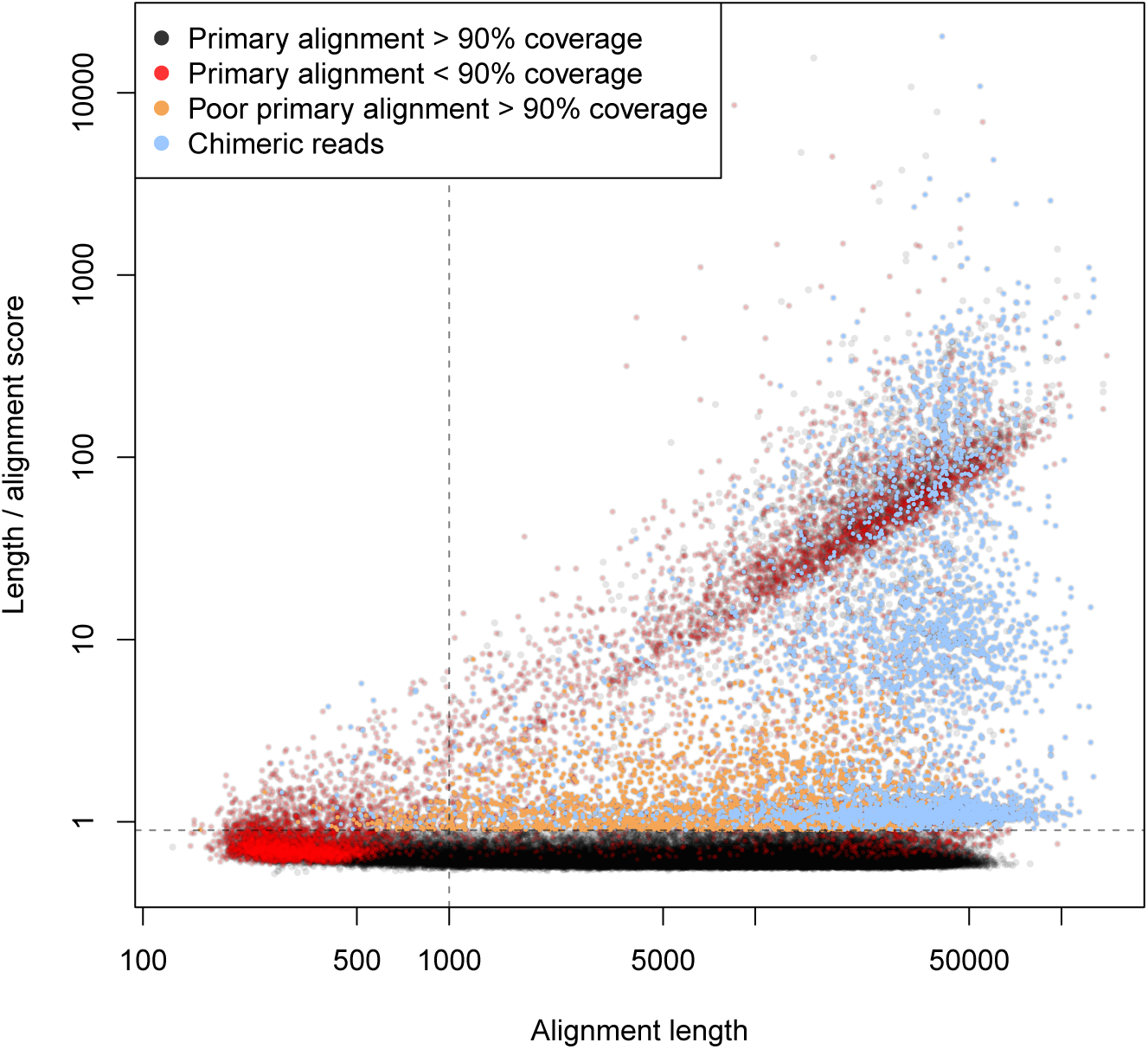
Relationship of read length and alignment score for *Algoriphagus alkaliphilus*. Chimeric reads (where both the negative and positive strand of a chromosomal region are sequenced on the same read) are shown in blue. Reads above 1kb and below the dotted line are retained as true alignments by the filtering process, requiring greater than 90 % query coverage. Axes are log scale.

Estimates of genome quality, percent completion and percent redundancy for each genome was calculated with Anvi’o v6.0 (Eren et al. 2015) and are shown with other relevant metrics in Table 1. All genomes achieved greater than 98% completion and less than 3% redundancy, except for the genome (UBA1547 sp002422915) predicted to belong to the recently proposed Candidate Phyla Radiation (CPR), which had completion at 85% and redundancy at 4%. The low completion estimate for the CPR genome is expected, as reported previously (Eren 2016).

**Table 1:** Taxonomy prediction was done on polished circularized genomes using `anvi-es timate-genome-taxonomy` in default mode (anvi’o version 6.0) Estimates of completeness and redundancy, and number of rRNA genes were obtained using `anvi-estimate-genome-completeness` and `anvi-display-contigs-stat`. The number of tRNA genes was esimated using `tRNAscan-SE 2.0`. Mean Illumina read coverages were calculated from filtered reads against the polished genomes using `samtools depth -a` (v 1.9). Mean Nanopore coverages were calculated by mapping filtered nanopore reads using `minimap2` (v 2.17), and `samtools depth -a` (v 1.9).

| Length (Mb) | % GC | %C | %R | Illumina coverage | Nanopore coverage | # rRNA genes | # tRNA genes | Taxonomy |
| --- | --- | --- | --- | --- | --- | --- | --- | --- |
| 3.14 | 62.8 | 98.59 | 0.00 | 392 | 193 | 2 | 48 | *Parvibaculum* |
| 4.47 | 64.4 | 100.00 | 2.82 | 402 | 94 | 2 | 49 | *Rhizobiaceae* |
| 3.73 | 63.9 | 100.00 | 0.00 | 68 | 32 | 2 | 46 | Unknown |
| 3.84 | 63.5 | 98.59 | 1.41 | 184 | 67 | 4 | 45 | *Blastomonas* |
| 3.74 | 72.4 | 100.00 | 0.00 | 121 | 39 | 4 | 63 | *UBA2363* |
| 5.16 | 42.6 | 98.59 | 0.00 | 180 | 97 | 6 | 40 | *Algoriphagus* |
| 3.98 | 66.9 | 100.00 | 1.41 | 41 | 16 | 4 | 51 | *Tabrizicola* |
| 3.96 | 71.9 | 98.59 | 2.82 | 125 | 57 | 2 | 53 | *UBA4742* |
| 4.68 | 66.1 | 100.00 | 2.82 | 57 | 17 | 4 | 52 | *Rhodobacteraceae* |
| 5.79 | 66.3 | 100.00 | 2.82 | 35 | 14 | 4 | 55 | *Aquimonas* |
| 0.79 | 51.0 | 85.92 | 4.23 | 88 | 36 | 2 | 40 | *UBA1547* |
| 3.06 | 65.9 | 100.00 | 0.00 | 1107 | 355 | 2 | 48 | *Brevundimonas* |
| 2.88 | 64.4 | 100.00 | 0.00 | 94 | 35 | 2 | 44 | *Oceanicaulis* |

Average genomic GC content varied widely, from 42% up to 72% (Table 1). Estimates of taxonomies were predicted using `anvi-estimate-genome-taxonomy`. One contig did not have high enough percent identity to provide a prediction (https://github.com/merenlab/anvio/issues/1328).

### Resolving genomic variants of high-coverage genomes

While visualizing the two most high-coverage genomes, *Parvibaculum* and *Brevundimonas*, it became apparent that there was a region in each genome where filtered Nanopore coverage dropped while the unfiltered Nanopore and Illumina coverage remained consistent. Upon further inspection of the alignments, it became apparent that a sub-population of the filtered reads supported the assembly, while the alignments for another population of reads (in this case the majority in both cases) stopped at a single base. The cause for this was a genomic variant that was supported by a majority of the long reads. In the case of *Parvibaculum*, there was a 35 kb deletion in the reference that was supported by a minority of reads, but once inserted, the majority of the reads supported. However, ultra-long reads that are correctly mapped support both of these variants, suggesting that there is indeed a minor variant that does not contain the 35 kb region, and major variant that does (Figure 3). Upon annotation with `prokka` (Seemann 2014), the deleted region contains a predicted nickel and cobalt resistance protein cnrA, a copper-resistance protein copB, and a type 3 IS family transposase, suggesting that a recent horizontal gene transfer event occured that may provide a loss or gain of function to this species.

**Figure 3:**
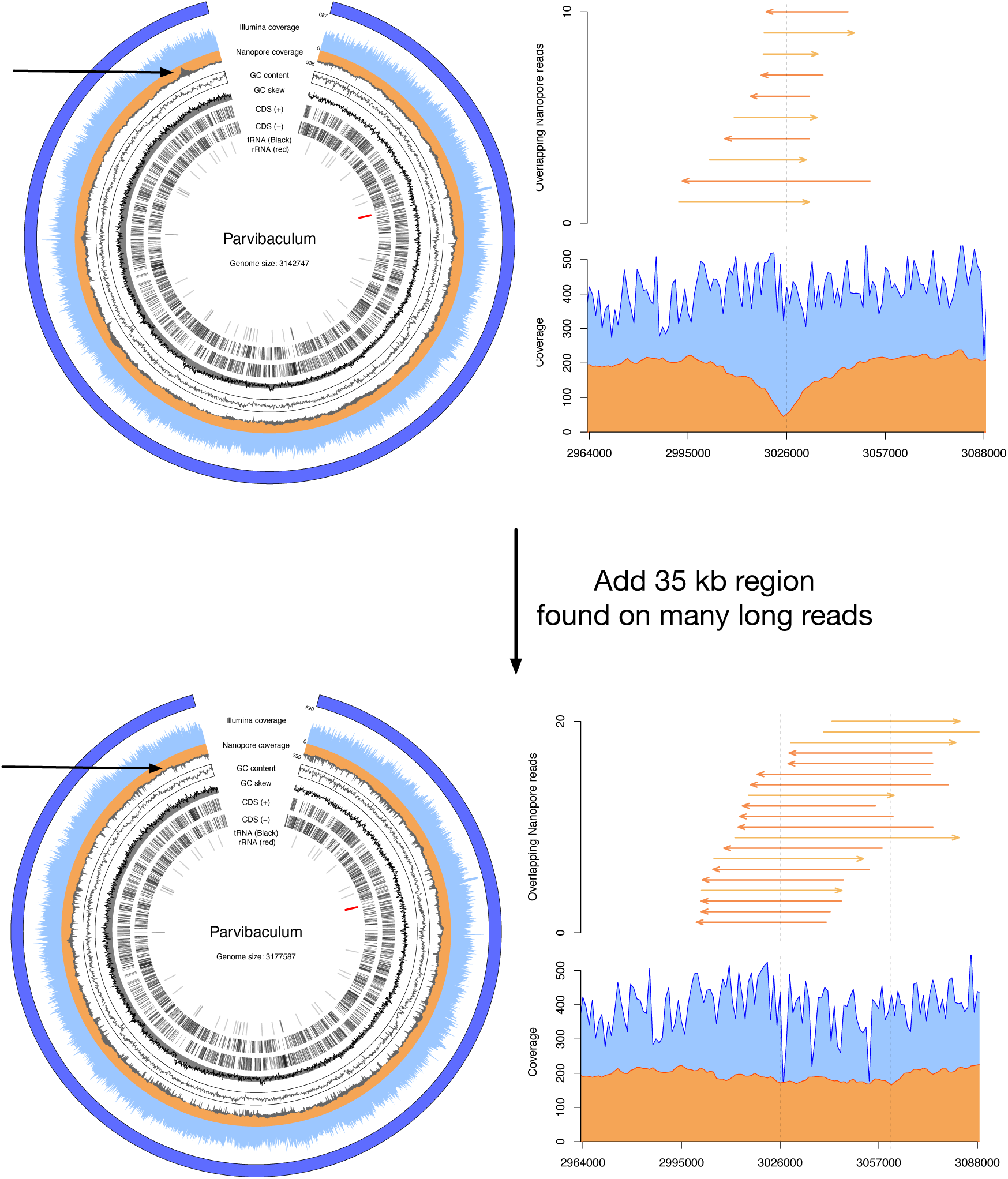
Visualization of the *Parvibaculum sp002480495* genome. The top half displays the initial assembly, and the bottom half with the addition of the 35 kb insertion. Arrows indicate the region of the insertion. The coverage for the region is shown in the right (Illumina blue, Nanopore orange), as well as reads overlapping the area with the deletion (reads mapping to positive strand: dark orange, reads mapping to negative strand: light orange). After inserting the region obtained from a read that completely spans the region, reads were mapped back to the assembly and plotted as before. Only a subset of reads were plotted as overlaps in both cases. Filtered Illumina and Nanopore coverage are shown in light blue and orange, respectively, unfiltered in grey. Coding sequencings (CDS), tRNAs, and rRNAs were predicted using prokka. Predicted rRNA genes are shown on inside track in red. Coverage was calculated using mosdepth in 1000 base windows. GC content and skew were calculated **8** with 1000 base windows. Light red regions (outer layer) highlights regions with less than 20 fold Illumina coverage.

### Genome of *UBA1547 sp002422915*

Visualization of the UBA1547 genome is shown in Supplemental figure 2. This genome is predicted to belong to the Candidate Phyla Radiation. There is an approximately 30 kb region (R1) where both Illumina and Nanopore coverage drop by 1.4X, and the GC content drastically increases (from 51% to 63-68%). While significant changes like these are often indicative of mis-assemblies or contaminant sequences in a metagenomic bin, overlapping long reads mapping to adjacent regions were found. There were 4 ultra-long reads that span the entire region, and also ultra-long reads (minimum 15kb) that overlap with junctions on either side (Figure 4). Annotation of this region revealed a type IV secretion system, a recombinase, and a type I endonuclease.

**Figure 4:**
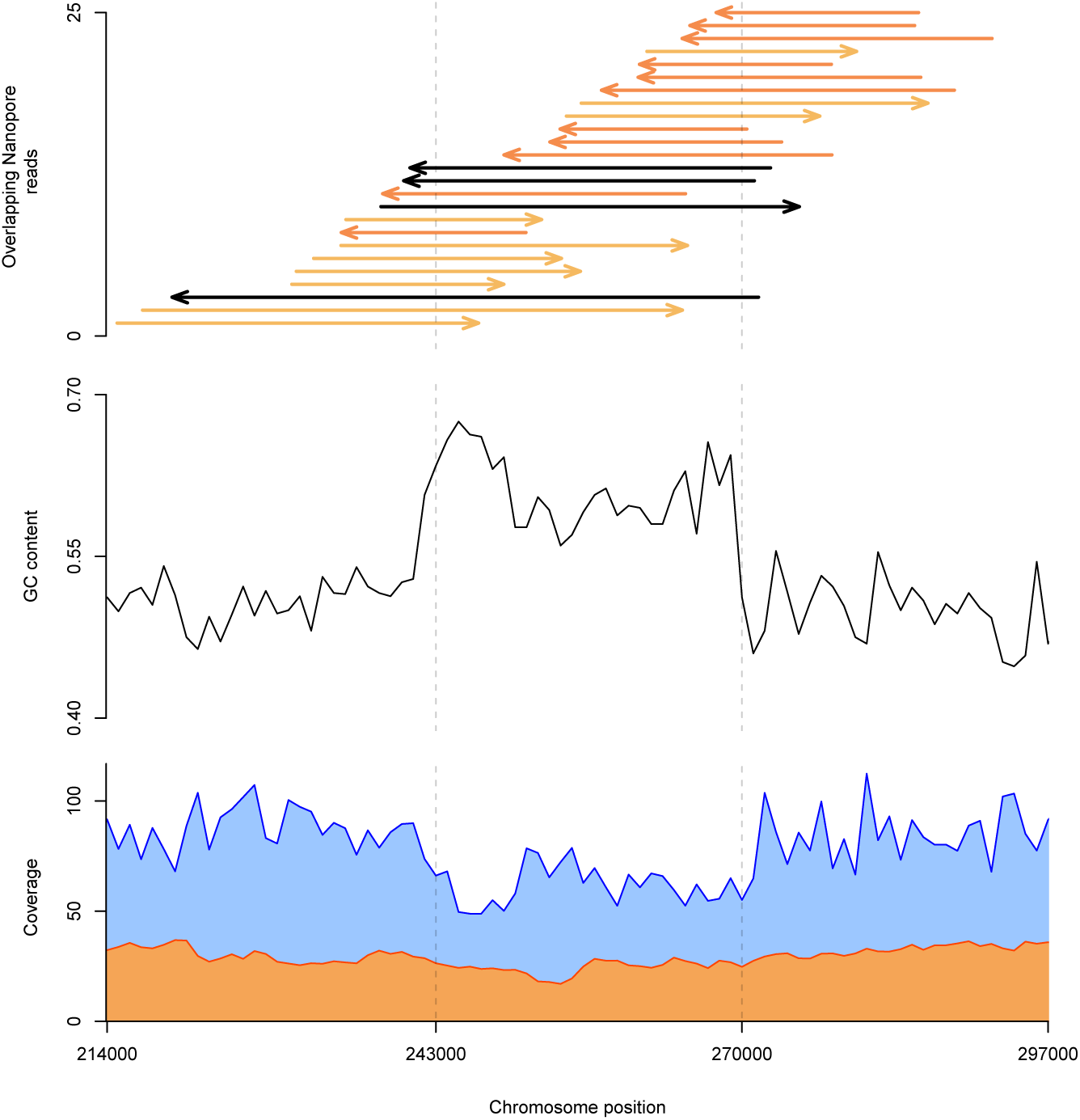
Deeper analysis of anomolous region in the UBA1547 sp002422915 genome. Top; Spanning and overlapping long reads at this region. Middle; GC content; Bottom; Illumina coverage (blue) and Nanopore coverage (orange). Only reads that have > 90% coverage, are greater than 15 kb, and overlap the junctions (243 kb and 270 kb) are shown. Reads that map to the positive strand are light orange, reads that map to the reverse strand are dark orange. Black reads completely span the region. GC content increases from about 51% to an average of 63% over the whole region. GC content and coverage calculated over 1000 base windows using mosdepth.

To ensure each genome is correctly assembled, visualizations of coverage, GC skew, and GC content are plotted for each genome (Supplemental figures 2-13). We observed that 63.77% of Nanopore reads above 1 kb and 64.38 % of uniquely mapped, properly paired Illumina reads are recruited by one of assembled genomes. Typically for isolate genome assemblies, paired-end short read information is used to detect potential mis-assemblies using a tool such as REAPR (Hunt et al. 2013). For metagenomic assemblies however, we observed that regions that were duplicated in either the genome or the population (e.g. duplicated 16S rRNA genes or transposases) will prevent paired end reads from uniquely mapping, causing REAPR to throw a false positive error. These reads must be removed as read support because we cannot uniquely map them to a single location. The result of this filtering may cause an apparent drop in Illumina coverage to zero. While we cannot accurately map Illumina reads to these regions, nanopore reads can still be used as support at these locations since they span the entire region or overlap with adjacent regions. For these cases, as long as Nanopore reads that spanned the entire region or overlapped with both adjacent regions, we accepted the region as correctly assembled.

### Are these genomes actually completed?

We re-assembled the filtered “psuedo-isolate” reads for each genome using alternative long read or hybrid assembly algorithms to determine if other assembly algorithms agree with the initial metagenomic assembly. Most pseudo-isolates succesfully re-assembled using using Flye in single genome mode, except for those with extremely low coverage (Table 2). Notably, Illumina-only Spades assembly performed poorly, and the hybrid assembler resulted in many contigs for very low coverage genomes. It may be important to note that a recent benchmarking of long-read assemblers suggests that Flye can assemble down to ~10X coverage and may be better at assembling low-coverage genomes than other algorithms (Ryan R Wick and Holt 2019). High coverage genomes successfully re-assembled with at least one other tool.

**Table 2:** Re-assembled genomes with “pseudo-isolates” long reads. Genome re-assembly was done using `Flye` v2.6 with parameters -g 5m, `Unicycler` v0.4.8 with parameters −1 −2 -l –no-pilon in hybrid mode, and `Unicyclyer` v0.4.8 with parameters-l –no-pilon in long-read only mode. UBA1547 and Rhodobacteraceae required using Flye in `--meta` mode to achieve circularization. The linear contigs did not re-circularize in `--meta` mode.

| Predicted Taxonomy | Flye Reassembled | Unicycler Hybrid Reassembled | Miniasm Reassembled | Spades Reassembled |
| --- | --- | --- | --- | --- |
| *Parvibaculum* | 1 circular contig | 1 circular contig | 17 linear contigs | 92 linear contigs |
| *Rhizobiaceae* | 1 circular contig | 1 circular contig | 4 linear contigs | 1866 linear contigs |
| Unknown | 1 circular contig | 1 circular contig | 1 linear contig | 258 linear contigs |
| *Blastomonas* | 1 circular contig | 2 linear contigs | 4 linear contigs | 135 linear contigs |
| *UBA2363* | 1 circular contig | 1 circular contig | 1 linear contig | 366 linear contigs |

Complete and validated genomes from a metagenome
| Predicted<br>Taxonomy | Flye<br>Reassembled | Unicycler<br>Hybrid<br>Reassembled | Miniasm<br>Reassembled | Spades<br>Reassembled |
| --- | --- | --- | --- | --- |
| *Algoriphagus* | 1 circular contig | 6 linear contigs | 1 linear contig | 417 linear contigs |
| *Tabrizicola* | 1 linear contig | 15 linear contigs | 4 linear contigs | 225 linear contigs |
| *UBA4742* | 1 circular contig | 1 circular contig | 4 linear contigs | 211 linear contigs |
| *Rhodobacteraceae* | 1 circular contig | 27 linear contigs | 3 linear contigs | 404 linear contigs |
| *Aquimonas* | 1 linear contig | 26 linear contigs | 1 linear contig | 1224 linear contigs |
| *UBA1547* | 1 circular contig | 1 circular contig | 1 linear contig | 5 linear contigs |
| *Brevundimonas* | 1 circular contig | 1 circular contig | 50 linear contigs | 39 linear contigs |
| *Oceanicaulis* | 1 linear contig | 1 circular contig | 4 linear contigs | 37 linear contigs |

## 0.5 Discussion

Here, we demonstrate that a recently published long-read metagenomic assembly algorithm metaFlye can be used to generate circularized genomes from an ultra-long read metagenomic dataset (N50 > 25 kb) when filtered and polished with paired-end Illumina reads. The genomes recruit 63.77 % of Nanopore reads and 64.38 % of Illumina reads. We also propose a general pipeline to visually validate *de novo* assembled genomes from metagenomic data when no reference genomes are available.

### Ultra-long reads are essential

We have observed that ultra-long sequencing reads are required to obtain highly contiguous genomes directly from metagenomic data using metaFlye. Ultra-long reads are needed to resolve repetitive DNA sequences and large indel strain variants, and can provide evidence that apparent anomalous DNA regions truly belong in the genome, and may be due to recent horizontal gene transfer. In the UBA1547 genome, we would anticipate that the short-read only metagenomic assembly and binning technique would completely miss the region with a drop in coverage and higher GC content because common binning algorithms rely on tetranucleotide frequency and coverage to cluster contigs into MAGs. There are several possibilities for explaining the GC content and coverage differences. Firstly, it could be that the coverage changes simply to due PCR bias in the Illumina library preparation step. Secondly, it could also be that this region is a self-excising mobile genetic element. No reads were found that support an excised DNA element, however, the coverage may be too low (35X) to be certain. Resequencing with higher coverage may provide better evidence for whether this is an artifact of Illumina sequence or an functional mobile genetic element. New algorithms using Oxford Nanopore’s ReadUntil API such as UNCALLED (Kovaka et al. 2020) can selectively enrich Nanopore reads in real-time, and may be useful for selectively resequencing this genome from the same DNA input.

### Validation requires accurate read mapping

When mapping reads back to metagenomic assemblies, it is important to remember that many genes in the community may be duplicated. Highly conserved genes such as the 16S rRNA gene can be problematic because multiple alignments with equal mapping quality may be reported for a pair of Illumina reads, making it impossible to identify where the read pair truly maps to. If short reads cannot be unambiguously mapped, they must be removed or they may cause errors when generating the consensus sequence for the genome.

The same is true for long reads, however it can be easier to resolve ambiguities with long reads by using only reads that anchor to adjacent regions. Using the default reporting parameters in minimap2, all possible alignments are reported, including secondary and supplemental alignments. Furthermore, multiply mapped and chimeric reads required additional filtering to remove. Others have used similar filtering strategies (Caceres et al. 2019), but we propose an additional criterion to help reduce the number of chimeric reads. When using the default scoring parameters of minimap2, we propose that a minimum read length of 1000 and a read length to alignment score ratio of less than 0.9 is used for filtering, in addition to percent query coverage of each read. For both the Illumina and Nanopore platforms, it is important to consider the limitations of the mapping algorithms used when validating a metagenomically-derived genome.

It is also important to highlight the need for an automated method to validate using long reads when short reads do not support the reference for technical reason. REAPR can be used on isolates, however, the assumption that each read pair will map uniquely breaks down in metagenomic datasets. It is therefore expected that false positives will be reported when using REAPR. To get around these, we manually evaluated long reads at problematic regions. An automated method could significantly improve the throughput, but further work is required for this.

### Genome variants can be resolved by visualizing long read support

Even though the initial assembly of *Parvibaculum* is supported by overlapping long reads, the majority of the population contains an insertion that could only be captured by visualizing filtered long reads. Ensuring that ultra-long reads are accurately mapped allowed us to resolve variants containing a mobile genetic element that may provide a functional advantage in the environment. Although a majority of reads support the insertion event, many reads (including ultra-long reads) do support a smaller population of *Parvibaculum* that does not contain the region. Taking this approach with ultra-long reads may improve the accuracy of genome assemblies and allow for the identification of recently horizontal gene transfer events or strain variants directly from metagenomes.

This highlights the need to visually validate each genome assembly obtained from metagenomic data. Even though unfiltered read coverage may support the reference, there may be anamolous regions that require filtering the reads to detect. Identification of these regions can allow for manual curation to completion (Figure 3).

### Limitations

This sequenced community was derived from an enrichment culture, which may be less diverse than an environmental sample or a gut microbiome sample. To achieve sequencing for more diverse samples, higher read depth may be needed. We only achieved approximately 7 gigabases of sequencing output from this flow cell; it is now typical to achieve twice that on a single R9.4.1 flow cell. Furthermore, the newer R10.3 flow cells may enable better consensus sequences.

This community does not have known reference genomes. It is therefore very difficult to gauge the genome quality using standard tools. REAPR can be used to estimate the number of bases that are supported by mapped Illumina reads, however, the number would be artificially decreased due to apparent zero coverage at repetitive regions. Given the recent developments in both Illumina and Oxford Nanopore platforms, a tool that evaluates the quality of *de novo* genome assemblies without the need for a reference would significantly benefit the metagenomic sequencing community.

It is our observation that the assembly of genomes is critically dependent on achieving ultra-long reads with a read N50 > 20 kb. Importantly, bead-beating and column based methods may not provide DNA fragments long enough to achieve circularization directly from the metagenomic data. Furthermore, in some environmental DNA samples, such as tailings ponds or soil samples, it may be more difficult to extract high quality high molecular weight DNA. Tough-to-lyse bacteria may be more resistant enzymatic lysis methods, so interpreting the relative abundance of the bacteria in these datasets may not be truly representative of the environment they were derived from. It is also worth noting that there is a large difference in the relative abundance of several genomes if calculated with Nanopore or Illumina reads, even though the input DNA for both runs was from the same extraction. More work is needed, but it is important to highlight that different platforms may result in different apparent relative abundances for bacteria that have highly diverse genomic characteristics.

### Future directions

For this experiment, we focused only on generating circular contigs > 300 kb to remove a mitochondrial genome that originated from algae in the population. In the future, we will further investigate this dataset to characterize circularized plasmids and other genomic elements of interest.

It is somewhat difficult to gauge exactly how many errors there are in the assemblies without known reference genomes. Furthermore, the more recent benchmarking for Nanopore consensus sequences with R9.4.1 flow cells suggests that typical consensus sequences of 100X coverage can return Q-scores of around 30. We found that many small indel corrections were necessary at homopolymeric regions, something that the R9.4.1 flow cell struggle with. The recently released dual pore head R10.3 flowcell is anticipated to better handle homopolymeric regions, so it will be required in future investigations to benchmark the R10.3 vs. R9.4.1 as it becomes available to the public.

## 0.6 Acknowledgements

We’d like to thank Suncor Energy for funding a portion of D.G.’s salary throughout this work. Document created with BiocStyle (Oleś, Morgan, and Huber 2020).

## 0.7 Data and code availability

Reads are available on the European Nucleotide Archive under project PRJEB36155. Code to reproduce the analysis and circos plots is available here https://doi.org/10.5281/zenodo.3745531.

## 0.8 Funding

Suncor Energy

NSERC RGPIN-03878-2015 (GBG)

NSERC RGPIN-05214-2015 (LMG)

CIHR PJT-159708 (GBG)

Ontario Graduate Scholarship (DJG)

