## supplementary_materials.pdf for "Complete and validated genomes from a metagenome"

**03 April 2020**

### 1 Supplementary materials

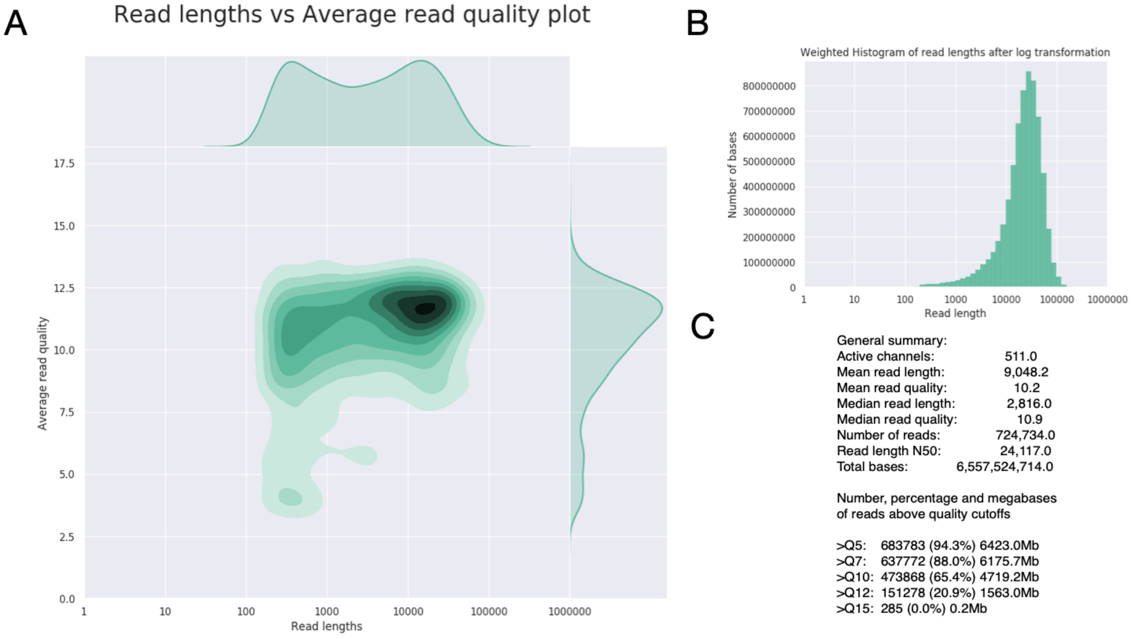


**Figure 1: Read length statistics of Nanopore run**
NanoPlot (Coster et al., 2018) was used to calculate statistics directly from raw Guppy 3.3.0 output. A, Read lengths vs read quality. B, Histogram of read length by bases obtained. C, Select stats obtained from NanoStats.txt

**Supplementary Table 1: Read recruitment of polished genomes.** Percentage of reads mapped was calculated from the filtered subset of reads for both Nanopore and Illumina. For Nanopore, filtering was performed as described in methods. Raw reads below 1 kb bases were removed before counting total number of Nanopore reads (final number is 440,931) since this no reads below 1 kb were considered for mapping. Number of reads for Illumina was counting after quality control and trimming (216,924,142 in total).

| Predicted taxonomy | Percent of Nanopore reads recruited | Percent of Illumina reads recruited | % GC |
|---|---|---|---|
| Parvibaculum | 13.96 | 7.75 | 62.8 |
| Rhizobiaceae | 5.95 | 11.30 | 64.4 |
| Unknown | 2.37 | 1.59 | 63.9 |
| Blastomonas | 4.14 | 4.43 | 63.5 |
| UBA2363 | 2.31 | 2.87 | 72.4 |
| Algoriphagus | 8.81 | 5.88 | 42.6 |
| Tabrizicola | 1.04 | 1.04 | 66.9 |
| UBA4742 | 3.62 | 3.12 | 71.9 |
| Rhodobacteraceae | 1.23 | 1.68 | 66.1 |
| Aquimonas | 1.55 | 1.29 | 66.3 |
| UBA1547 | 0.60 | 0.44 | 51.0 |
| Brevundimonas | 16.37 | 21.29 | 65.9 |
| Oceanicaulis | 1.82 | 1.70 | 64.4 |

### Supplementary materials

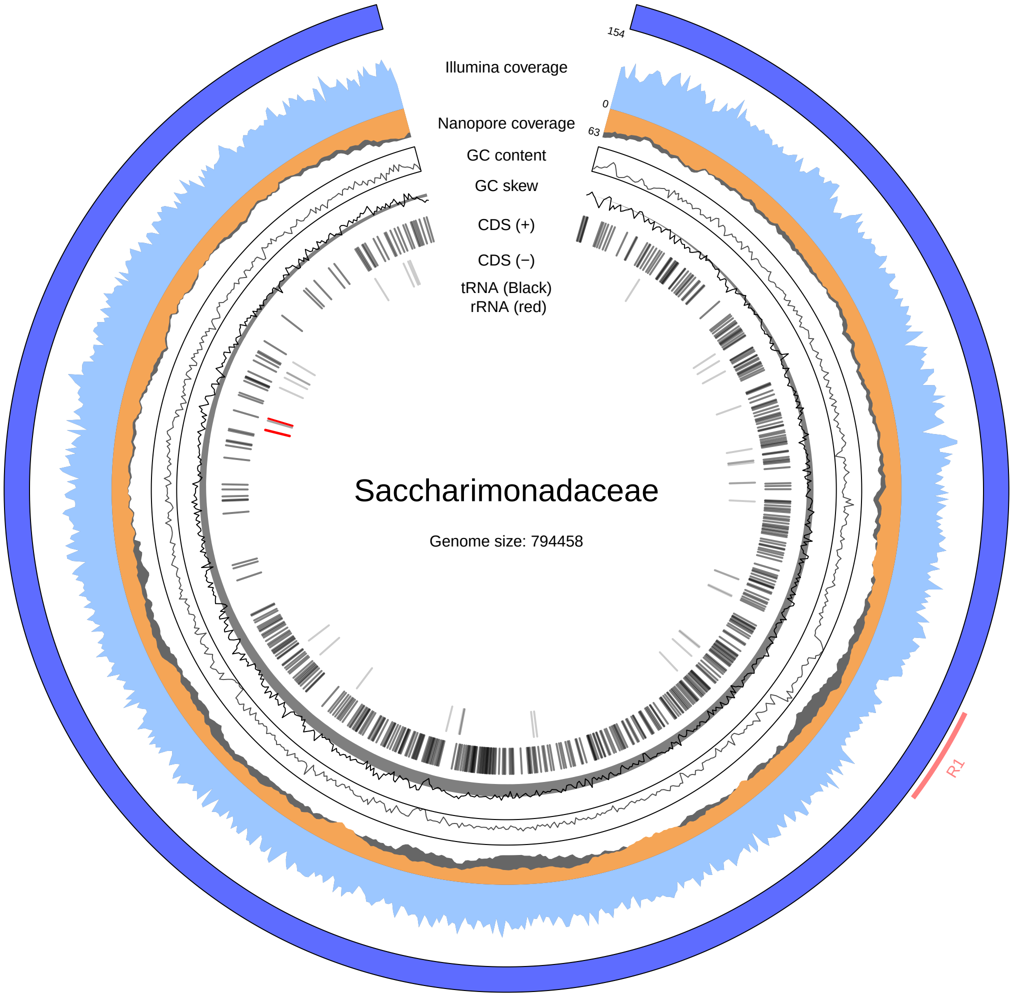


**Figure 2:** **Visualization of the UBA1547 sp002422915 genome**
Taxonomic prediction reveals Saccharimonadaceae, a genus belonging to the recently proposed Candidate Phylum Radation. Filtered Illumina and Nanopore coverage are shown in light blue and orange, respectively, unfiltered in grey. GC content ranges from 40 % to 67 %, with outward from the center increasing. Coding sequencings (CDS), tRNAs, and rRNAs were predicted using prokka. Predicted rRNA genes are shown on inside track in red. Coverage was calculated using mosdepth in 1000 base windows. GC content and skew were calculated with 1000 base windows. R1 (light red - outer layer) highlights the anamolous region.

### Supplementary materials

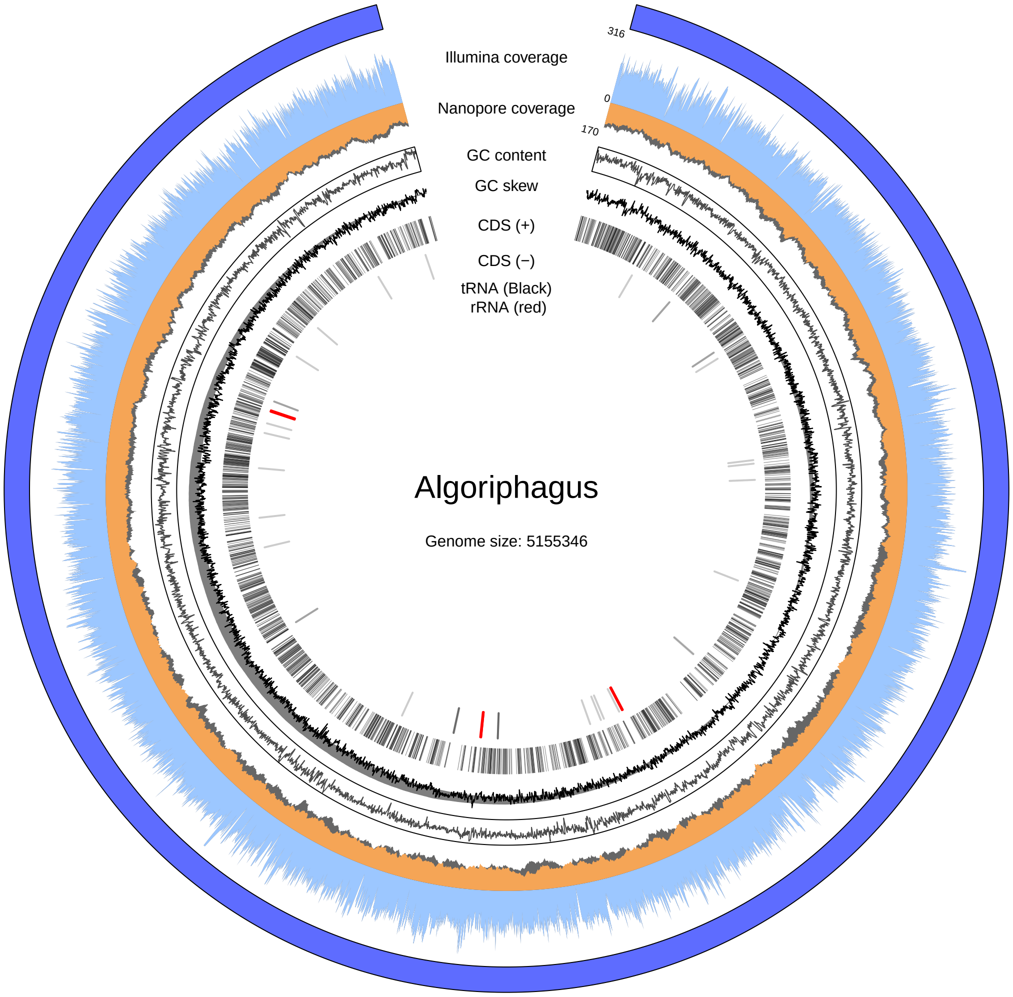


**Figure 3: Visualization of the Algoriphagus alkaliphilus genome**
Filtered Illumina and Nanopore coverage are shown in light blue and orange, respectively, unfiltered in grey. Coding sequencings (CDS), tRNAs, and rRNAs were predicted using prokka. Predicted rRNA genes are shown on inside track in red. Coverage was calculated using mosdepth in 1000 base windows. GC content and skew were calculated with 1000 base windows. Light red regions (outer layer) highlights regions with less than 20 fold Illumina coverage.

### Supplementary materials

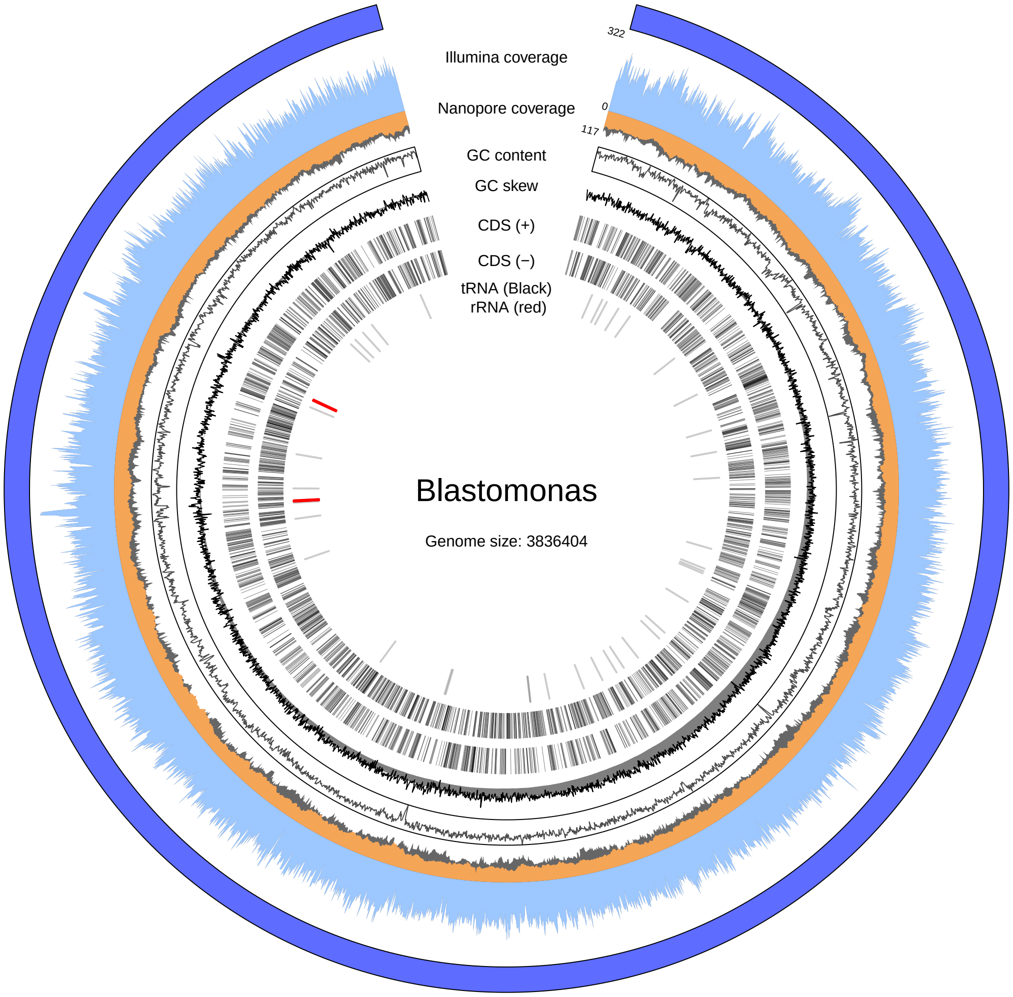


**Figure 4: Visualization of the Blastomonas sp001713435 genome**
Filtered Illumina and Nanopore coverage are shown in light blue and orange, respectively, unfiltered in grey. Coding sequencings (CDS), tRNAs, and rRNAs were predicted using prokka. Predicted rRNA genes are shown on inside track in red. Coverage was calculated using mosdepth in 1000 base windows. GC content and skew were calculated with 1000 base windows. Light red regions (outer layer) highlights regions with less than 20 fold Illumina coverage.

### Supplementary materials

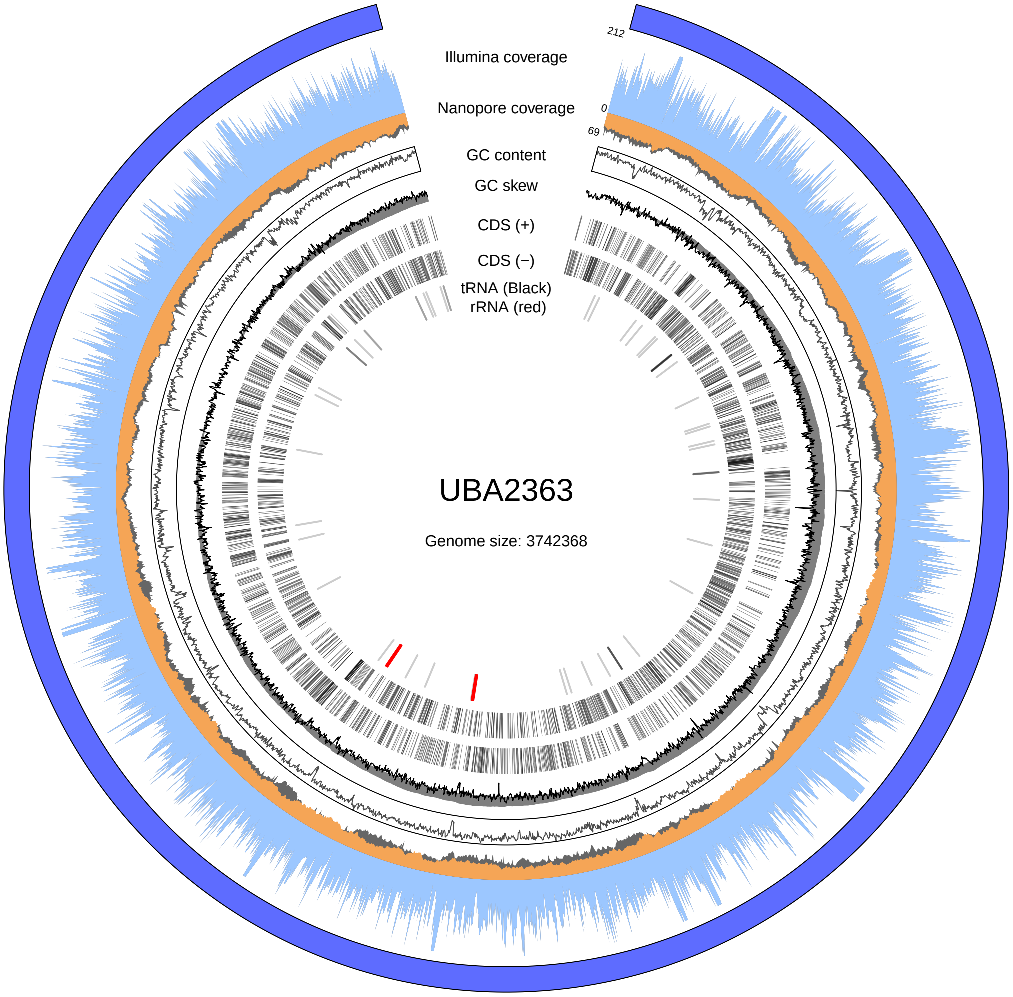


**Figure 5: Visualization of the UBA2363 sp002344355 genome**
Filtered Illumina and Nanopore coverage are shown in light blue and orange, respectively, unfiltered in grey. Coding sequencings (CDS), tRNAs, and rRNAs were predicted using prokka. Predicted rRNA genes are shown on inside track in red. Coverage was calculated using mosdepth in 1000 base windows. GC content and skew were calculated with 1000 base windows. Light red regions (outer layer) highlights regions with less than 20 fold Illumina coverage.

### Supplementary materials

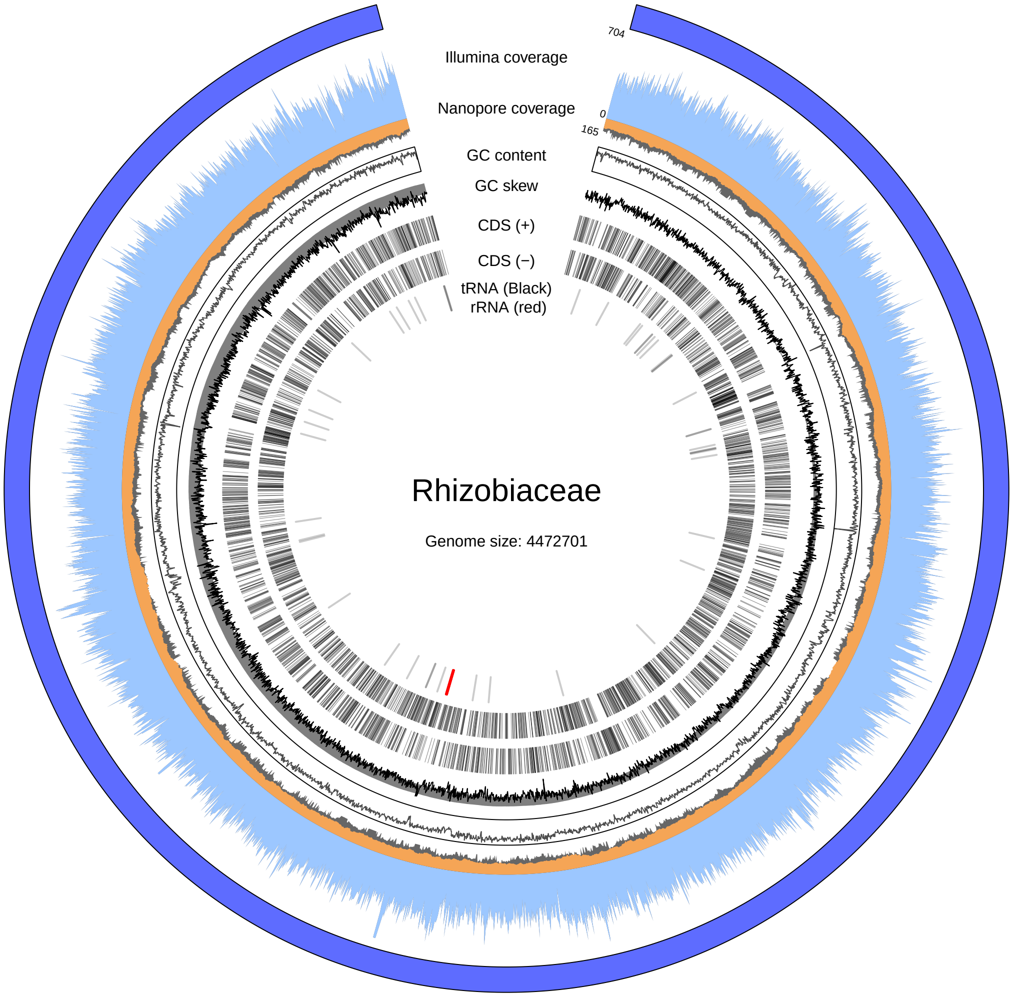


**Figure 6:** **Visualization of the Rhizobiaceae genome**
Filtered Illumina and Nanopore coverage are shown in light blue and orange, respectively, unfiltered in grey. Coding sequencings (CDS), tRNAs, and rRNAs were predicted using prokka. Predicted rRNA genes are shown on inside track in red. Coverage was calculated using mosdepth in 1000 base windows. GC content and skew were calculated with 1000 base windows. Light red regions (outer layer) highlights regions with less than 20 fold Illumina coverage.

### Supplementary materials

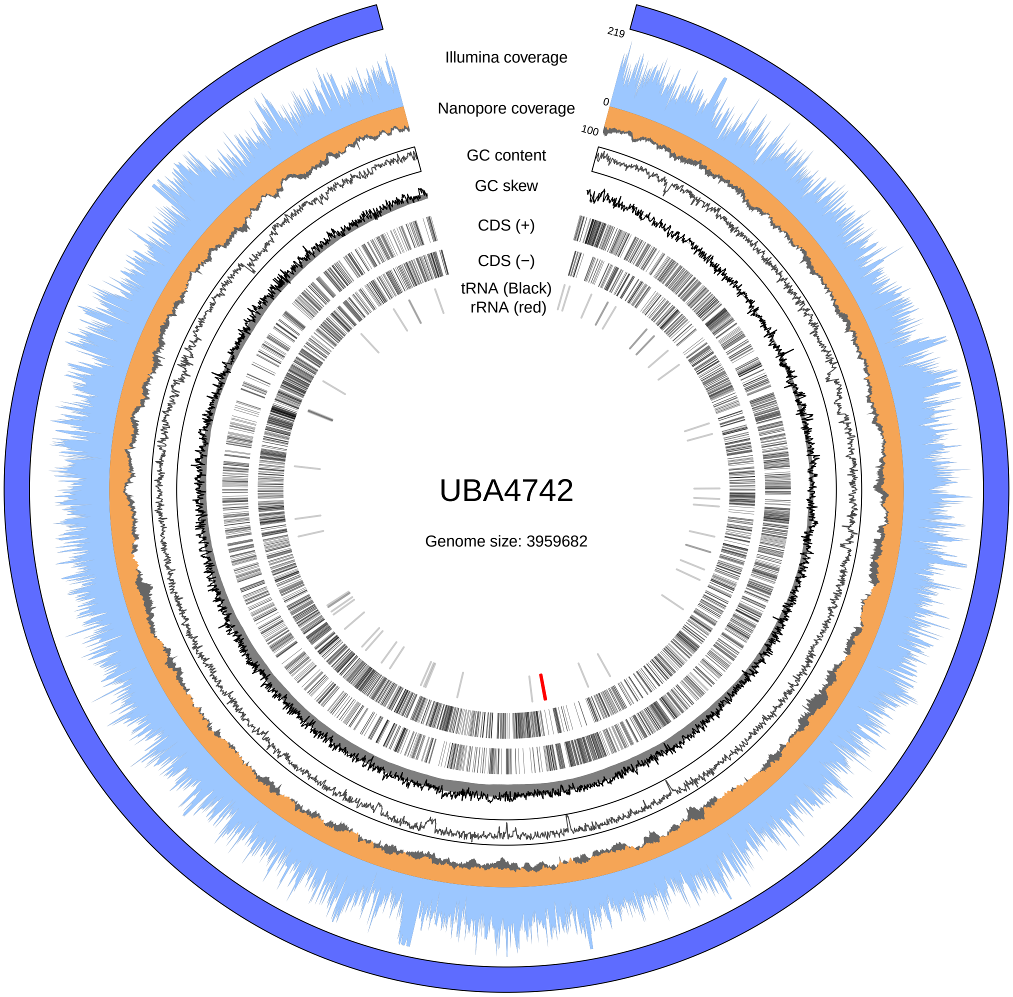


**Figure 7:** **Visualization of the UBA4742 sp002403895 genome**
Filtered Illumina and Nanopore coverage are shown in light blue and orange, respectively, unfiltered in grey. Coding sequencings (CDS), tRNAs, and rRNAs were predicted using prokka. Predicted rRNA genes are shown on inside track in red. Coverage was calculated using mosdepth in 1000 base windows. GC content and skew were calculated with 1000 base windows. Light red regions (outer layer) highlights regions with less than 20 fold Illumina coverage.

### Supplementary materials

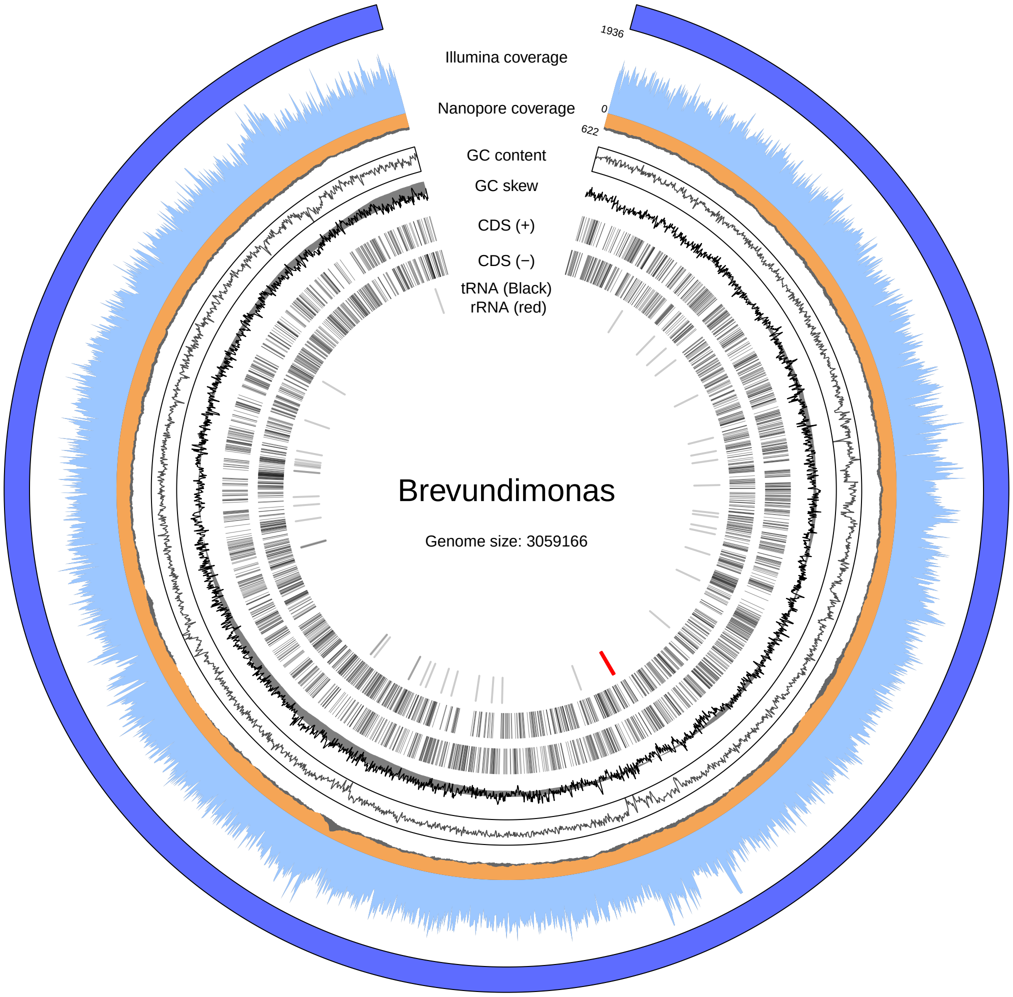


**Figure 8:** **Visualization of the Brevundimonas genome**
Filtered Illumina and Nanopore coverage are shown in light blue and orange, respectively, unfiltered in grey. Coding sequencings (CDS), tRNAs, and rRNAs were predicted using prokka. Predicted rRNA genes are shown on inside track in red. Coverage was calculated using mosdepth in 1000 base windows. GC content and skew were calculated with 1000 base windows. Light red regions (outer layer) highlights regions with less than 20 fold Illumina coverage.

### Supplementary materials

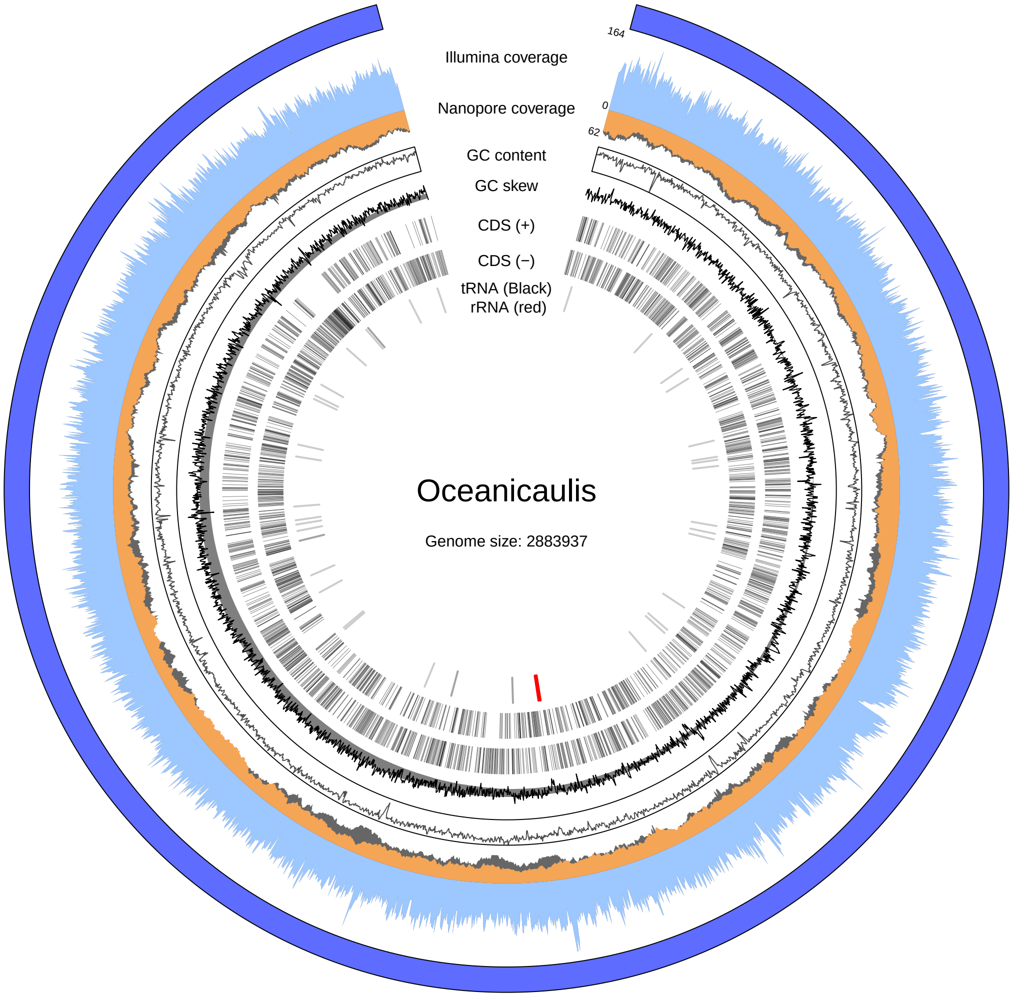


**Figure 9: Visualization of the Oceanicaulis sp000744995 genome**
Filtered Illumina and Nanopore coverage are shown in light blue and orange, respectively, unfiltered in grey. Coding sequencings (CDS), tRNAs, and rRNAs were predicted using prokka. Predicted rRNA genes are shown on inside track in red. Coverage was calculated using mosdepth in 1000 base windows. GC content and skew were calculated with 1000 base windows. Light red regions (outer layer) highlights regions with less than 20 fold Illumina coverage.

### Supplementary materials

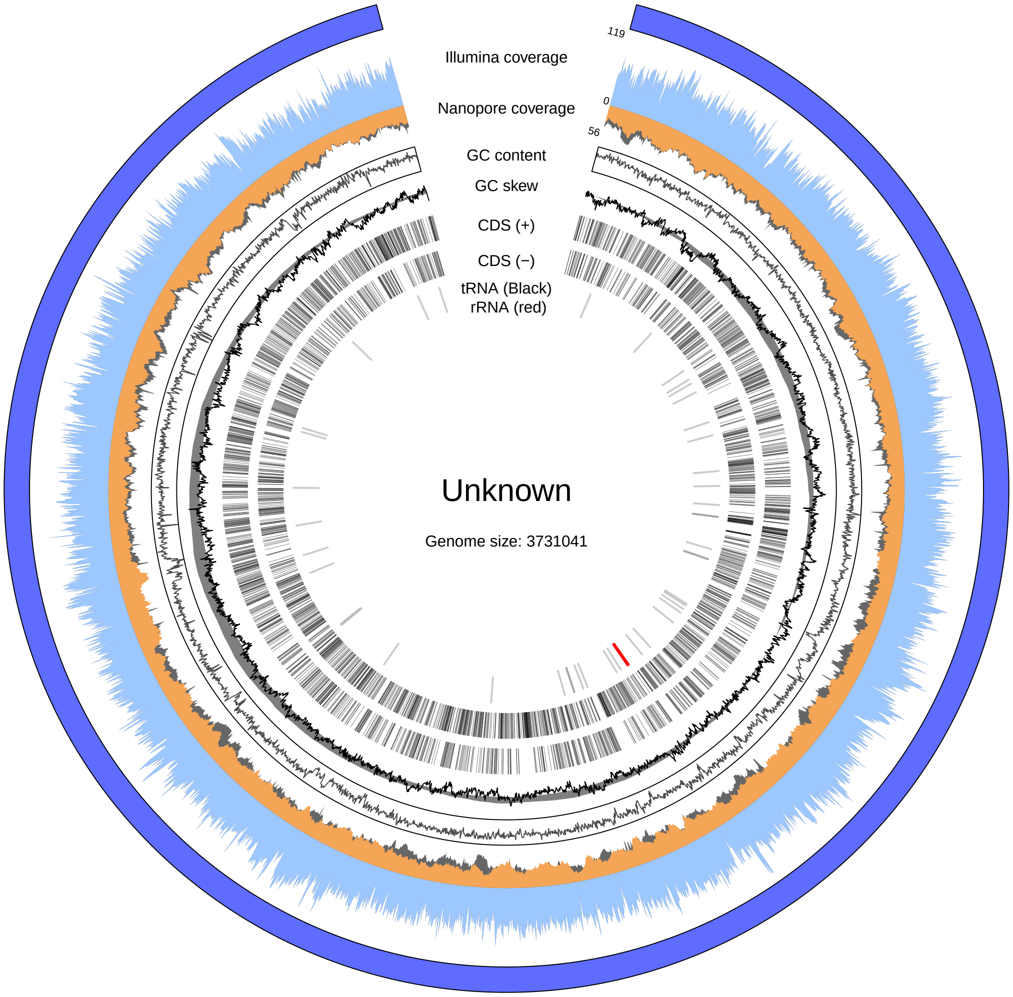


**Figure 10: Visualization of the Unknown genome**
Filtered Illumina and Nanopore coverage are shown in light blue and orange, respectively, unfiltered in grey. Coding sequencings (CDS), tRNAs, and rRNAs were predicted using prokka. Predicted rRNA genes are shown on inside track in red. Coverage was calculated using mosdepth in 1000 base windows. GC content and skew were calculated with 1000 base windows. Light red regions (outer layer) highlights regions with less than 20 fold Illumina coverage.

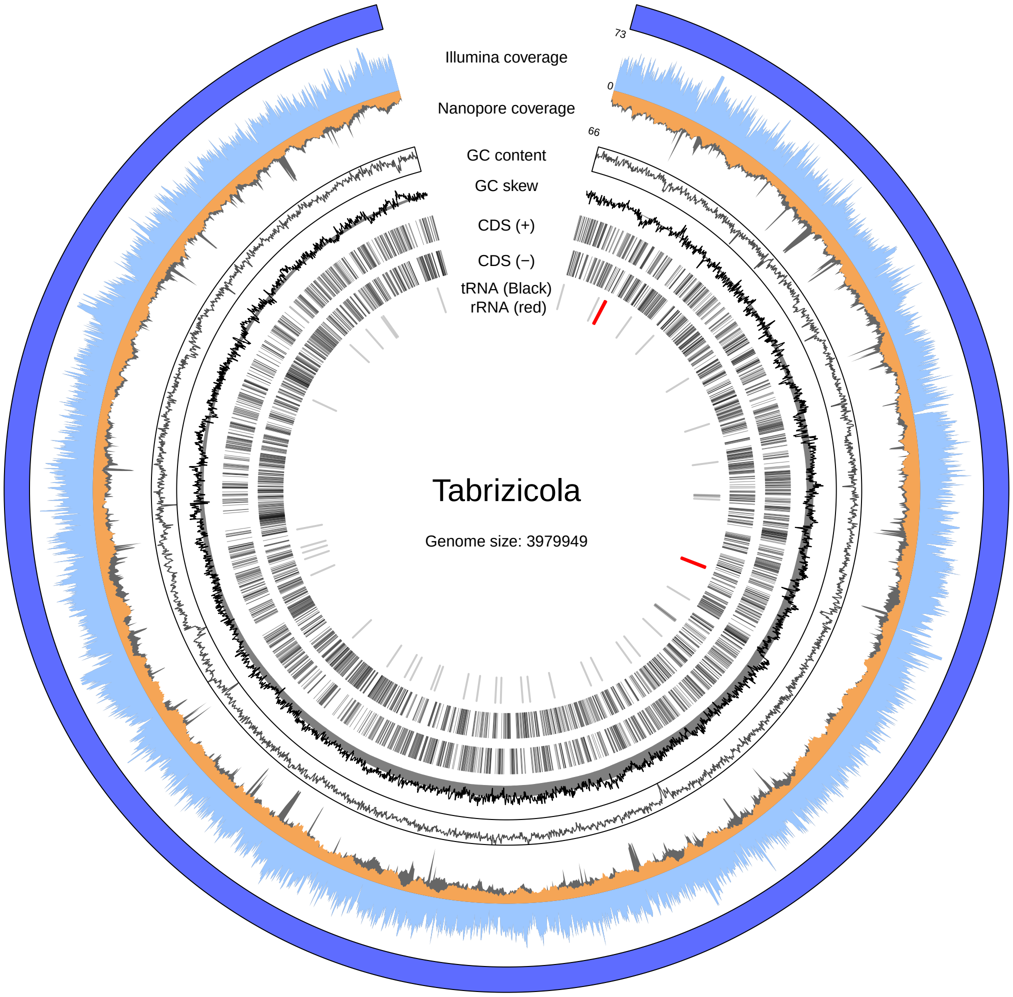


**Figure 11:** **Visualization of the Tabrizicola aquatica genome**
Filtered Illumina and Nanopore coverage are shown in light blue and orange, respectively, unfiltered in grey. Coding sequencings (CDS), tRNAs, and rRNAs were predicted using prokka. Predicted rRNA genes are shown on inside track in red. Coverage was calculated using mosdepth in 1000 base windows. GC content and skew were calculated with 1000 base windows. Light red regions (outer layer) highlights regions with less than 20 fold Illumina coverage.

### Supplementary materials

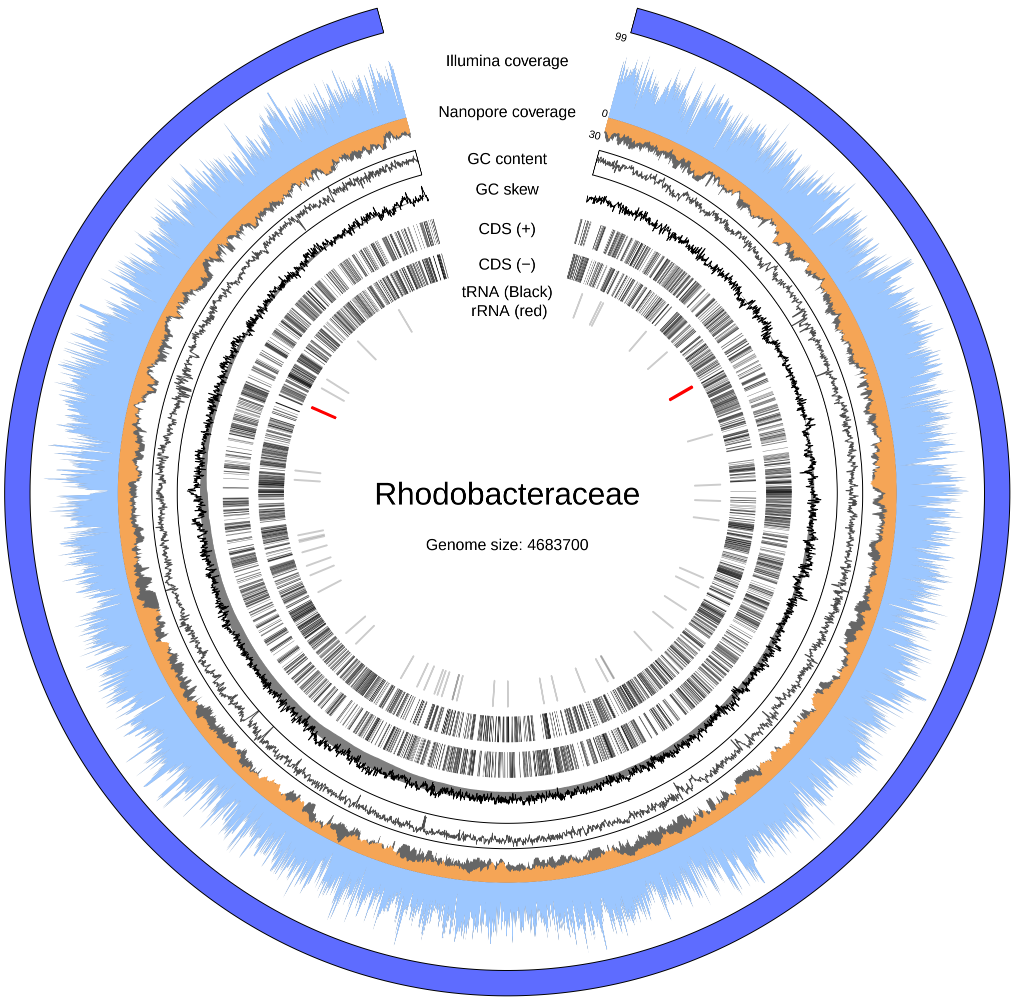


**Figure 12: Visualization of the Rhodobacteraceae genome**
Filtered Illumina and Nanopore coverage are shown in light blue and orange, respectively, unfiltered in grey. Coding sequencings (CDS), tRNAs, and rRNAs were predicted using prokka. Predicted rRNA genes are shown on inside track in red. Coverage was calculated using mosdepth in 1000 base windows. GC content and skew were calculated with 1000 base windows. Light red regions (outer layer) highlights regions with less than 20 fold Illumina coverage.

### Supplementary materials

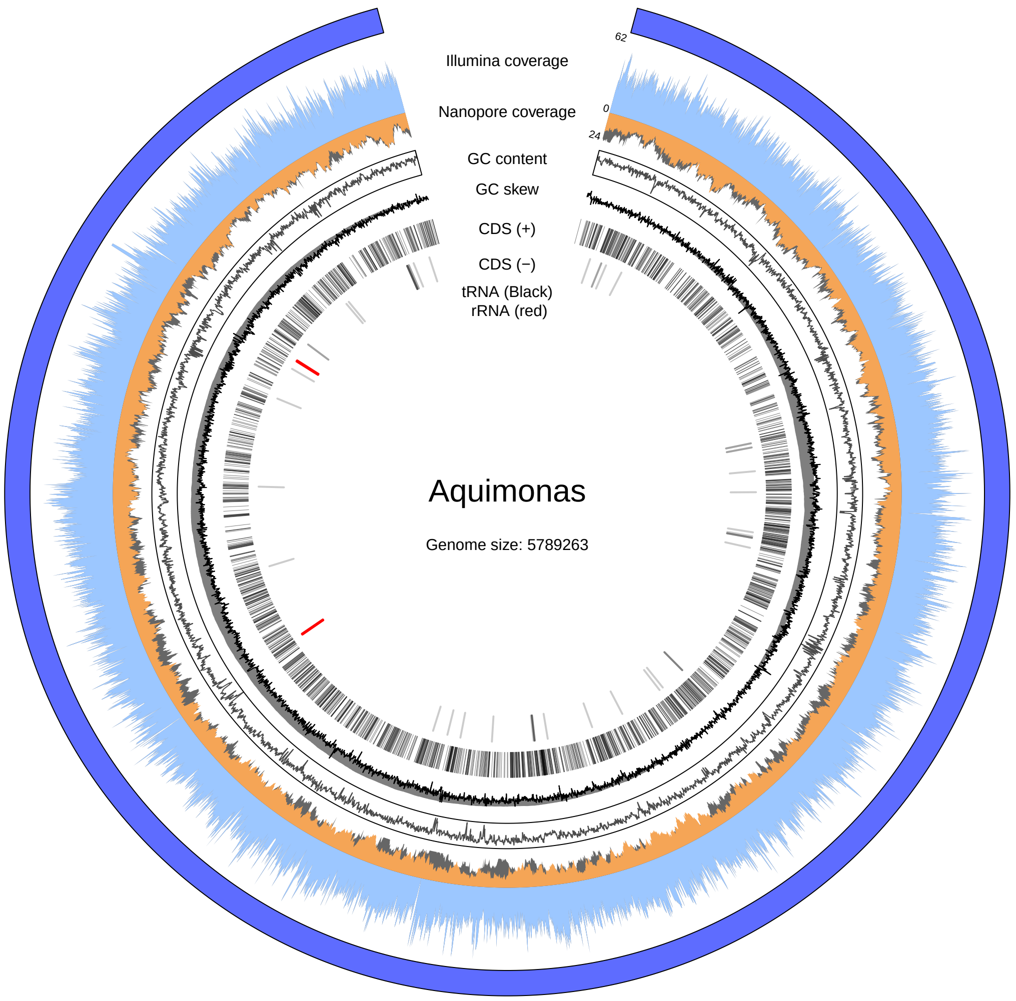


**Figure 13:** **Visualization of the Aquimonas voraii genome**
Filtered Illumina and Nanopore coverage are shown in light blue and orange, respectively, unfiltered in grey. Coding sequencings (CDS), tRNAs, and rRNAs were predicted using prokka. Predicted rRNA genes are shown on inside track in red. Coverage was calculated using mosdepth in 1000 base windows. GC content and skew were calculated with 1000 base windows. Light red regions (outer layer) highlights regions with less than 20 fold Illumina coverage.
